# Deep convolutional neural networks for predicting the quality of single protein structural models

**DOI:** 10.1101/590620

**Authors:** Jie Hou, Renzhi Cao, Jianlin Cheng

## Abstract

Predicting the global quality and local (residual-specific) quality of a single protein structural model is important for protein structure prediction and application. In this work, we developed a deep one-dimensional convolutional neural network (1DCNN) that predicts the absolute local quality of a single protein model as well as two 1DCNNs to predict both local and global quality simultaneously through a novel multi-task learning framework. The networks accept sequential and structural features (i.e. amino acid sequence, agreement of secondary structure and solvent accessibilities, residual disorder properties and Rosetta energies) of a protein model of any size as input to predict its quality, which is different from existing methods using a fixed number of hand-crafted features as input. Our three methods (InteractQA-net, JointQA-net and LocalQA-net) were trained on the structural models of the single-domain protein targets of CASP8, 9, 10 and evaluated on the models of CASP11 and CASP12 targets. The results show that the performance of our deep learning methods is comparable to the state-of-the-art quality assessment methods. Our study also demonstrates that combining local and global quality predictions together improves the global quality prediction accuracy. The source code and executable of our methods are available at: https://github.com/multicom-toolbox/DeepCovQA

## Introduction

In the past few decades, protein structure prediction had achieved significant progress on both template-based modeling and template-free modeling^1-7^. As a quality control step of modeling, protein model quality assessment (QA) plays an important role in selecting most accurate models among a massive number of decoys generated by protein structure modeling methods. There are two kinds of model quality assessment methods: local quality assessment^8-12^ and global quality assessment^10,13-22^. Local QA methods attempt to predict the spatial deviation of each residue in a model from the native structure (e.g. the absolute distance between the position of Ca atom of a residue in a model and that in the native structure), while global QA methods aim to predict the overall similarity (e.g. GDT-TS score^23^) between a model and its native structure. One kind of QA methods require a pool of models as input, which are often called consensus (or multi-model) methods^16,24-26^. Consensus QA methods evaluate a protein model by comparing it against the other models in the pool and calculating the average structural similarity as an indicator of the quality. Another kind of QA method only takes a single model as input to predict its quality, which are called single-model QA methods. These methods utilize the sequence and structural information of a single model itself to assess its quality. Consensus QA methods usually achieve good performance if a significant portion of models in the model pool are of good quality. However, it tends to fail if most models are of poor quality and is time-consuming if the size of the model pool is large. In contrast, the performance of single-model QA methods can be more consistent and more independent of the distribution of model quality in a pool because it predicts the quality of a model using only the information about itself. This is particularly useful if there are very few good models in a large model pool, which often happens in template-free protein structure prediction. Recent top-ranked single-model QA methods generally start with generating structure-related features from a model followed by applying machine learning methods to estimate its local or global quality score. Several features have been proved to be effective, such as sequence/profile alignment, predicted secondary structure and solvent accessibility of residues^8^, residue-residue contact potential^19^, torsion angle of main chain^27^, physicochemical properties^13^, and energy-based environment of residues and models^12,13,15^. Methods such as support vector machine^8,15,28^, neural network^14^, and linear combination ^22,29^ are commonly used for quality estimation. Many top QA methods have been largely tested and assessed in the Critical Assessment of Techniques for Protein Structure Prediction (CASP)^9^. ProQ2^10^ had the good performance on local quality assessment by using machine learning on the features including secondary structure, surface area, contacts information and so on, and its new version ProQ3^15^ that added Rosetta energies as features further improved quality assessment. DeepQA^14^ integrated energy-based potential scores with other structural information (i.e. RWplus^30^, OPUS^31^ and DFIRE^32^) derived from structures and improved the global quality prediction. Qprob^13^ combined structural/sequence features, including energy and physicochemical properties of a model, to evaluate its quality. All the methods predict local or global quality separately. No methods tried to predict both quality measurements at the same time, even though some methods derived the global score converted from predicted local quality score of residues^8,33^. Moreover, traditional machine learning based quality assessment methods used a fixed-size sliding window approach to estimate the local deviation of each residue, in which the features of neighboring residues within a window of a specific size (e.g. 5, 11 and 21 residues) that is centered on a target residue are combined by machine learning approaches to predict the local quality of the residue. Recently, deep learning techniques that can handle input of varied size have achieved significant success in the bioinformatics field^14,34,35^. Especially, the application of deep convolutional neural network (CNN)^7,36,37^ (e.g. 1DCNN for sequential data and 2DCNN for image-like inputs) has achieved the promising performance and becomes one of the best machine learning methods for solving bioinformatics problems^7,36,37^. The convolutional neural networks can learn longer-range sequential and structural information from the input features of arbitrary length, which cannot be utilized traditional sliding window approaches.

In this study, we designed novel deep convolutional networks to predict the local and global quality of a protein model consisting of any number of residues, leveraging their capability of handling input of any length. Furthermore, we used a novel multi-task learning framework to study whether global and local quality predictions can synergistically interact to improve prediction performance. Specifically, we developed three novel single-model predictors, InteractQA-net, JointQA-net and LocalQA-net, which use sequence information, structural features, residue-specific Rosetta energies, and other energy scores as input to predict local quality or both global and local quality of a model. We also combined the three predictors to further improve prediction accuracy.

## Result and Discussion

### Training and parameter optimization

We trained each convolutional network with different parameter setting on our training dataset and selected the best trained model using the ASE metric calculated on the validation dataset. We optimized the following hyper-parameters: the depth of the network (from 5 to 10), the filter size of each convolution layer (from 5 to 15), and the number of filters in each convolutional layer (from 5 to 20). Based on the results on the validation set, the depth of convolutional layers in the InteractQA-net is set to 10, number of filters to 10, and the filter size to 15. For the JointQA-net and LocalQA-net, the final depth of convolution layers is set to 5, number of filters to 5, and filter size to 6 in each convolutional layer. The deep networks trained with these parameters on the training dataset were evaluated on the independent test datasets.

### Comparison of local quality predictions with other single-model QA methods on CASP11 and CASP12

We compared InteractQA-net, JointQA and LocalQA-net with CASP single-model QA methods on the 1^*st*^ stage and 2^*nd*^ stage subsets of CASP11 and CASP12 test datasets. We calculated the average ASE score across all models of each subset for our three predictors and other CASP predictors for comparison (Table 1 and Table 2). LocalQA-net achieved slightly better performance than InteractQA-net and JointQA-net according to the average ASE scores on the CASP 11 datasets, but it was slightly worse than InteractQA-net and JointQA-net on the CASP 12 datasets, suggesting that including the global quality prediction did not necessarily help with the local quality prediction. However, the accuracy of the ensemble (CNNQA) of the three predictors is higher than each our three predictors, indicating that the three methods are complementary.

**Table 1:** The evaluation results (average ASE scores) of local quality predictions of single-model local quality QA predictors on stage 1 and stage 2 models from CASP 11.

| **Predictor** | **Stage1** | **Stage2** | **Average** |
| --- | --- | --- | --- |
| CNNQA | 81.06 | 78.43 | 79.75 |
| LocalQA-net | 79.90 | 78.26 | 79.08 |
| InteractQA-net | 80.27 | 76.97 | 78.62 |
| JointQA-net | 79.81 | 77.03 | 78.42 |
| Wang_deep_3 | 78.11 | 74.56 | 76.34 |
| Wang_deep_2 | 77.58 | 74.22 | 75.90 |
| ProQ2 | 75.87 | 75.74 | 75.81 |
| ProQ2-refine | 75.91 | 75.67 | 75.79 |
| Wang_deep_1 | 77.78 | 73.43 | 75.60 |
| Wang_SVM | 76.79 | 71.91 | 74.35 |
| MULTICOM-NOVEL | 67.13 | 67.11 | 67.12 |
| VoroMQA | 62.72 | 66.45 | 64.58 |
| MULTICOM-REFINE | 62.68 | 65.26 | 63.97 |
| MULTICOM-CLUSTER | 62.98 | 64.87 | 63.92 |
| MULTICOM-CONSTRUCT | 63.39 | 64.35 | 63.87 |

**Table 2:** The evaluation results (average ASE scores) of local quality predictions of single-model local quality QA predictors on stage 1 and stage 2 models from CASP12 datasets.

| Predictor | Stage1 | Stage2 | Average |
| --- | --- | --- | --- |
| CNNQA | 83.22 | 78.14 | 80.68 |
| ProQ3 | 82.19 | 78.54 | 80.37 |
| JointQA-net | 82.31 | 77.38 | 79.85 |
| InteractQA-net | 83.13 | 76.52 | 79.83 |
| LocalQA-net | 80.72 | 78.09 | 79.40 |
| Wang4 | 81.06 | 76.86 | 78.96 |
| Wang2 | 79.62 | 74.70 | 77.16 |
| VoroMQAsr | 79.32 | 74.69 | 77.01 |
| ProQ2 | 77.73 | 75.61 | 76.67 |
| VoroMQA | 78.87 | 74.26 | 76.56 |
| Wang1 | 72.22 | 72.52 | 72.37 |
| Wang3 | 53.86 | 60.55 | 57.20 |

The performance of LocalQA-net, InteractQA-net, and JointQA-net is comparable to the best performing predictors in CASP11 and CASP12 experiments (e.g. Wang_deep_3, ProQ2, and ProQ3), and CNNQA has slightly higher the average ASE score than all the CASP11 and CASP12 predictors.

### Comparison of global quality predictions with other single-model QA methods on CASP11 and CASP12

In order to evaluate the global quality prediction performance of our methods, we generated the global quality scores for our methods (InteractQA-net, LocalQA-net, LocalQA-net, CNNQA), which were converted from their residue-specific local quality predictions by averaging them using function 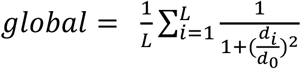. We compare them with other QA predictors on the same datasets of CASP 11 and CASP 12 used in the local quality prediction evaluation. We calculated the average Pearson’s correlation between predicted global quality scores and real global quality scores as well as the average loss to evaluate the performances of the QA predictors (see the results in Table 3 and Table 4). According to the Pearson’s correlation results on 1^*st*^ stage and 2^*nd*^ stage from CASP 11 and CASP 12, InteractQA-net achieved higher correlation and lower loss than LocalQA-net, which showed that integrating the global similarity into local quality prediction improved the global quality prediction derived from the local quality prediction. In terms of average Pearson’s correlation, InteractQA-net and CNNQA had the similar performance and both performed better than all other CASP11 and CASP12 predictors. In terms of average loss, InteractQA-net and CNNQA performed better than the other predictors on the CASP11 datasets, but worse than the two top methods (SVMQA^28^ and ProQ3^15^) on the CASP12 datasets.

**Table 3:** The evaluation results (Corr. – Pearson’s correlation and loss) of global quality predictions of single-model QA predictors on stage 1 and stage 2 models of CASP 11 datasets.

|  | **Stage1** |  | **Stage2** |  | **Average** |  |
| --- | --- | --- | --- | --- | --- | --- |
| **Feature** | **Corr** | **Loss** | **Corr** | **Loss** | **Corr** | **Loss** |
| InteractQA-net | 0.7243 | 0.0756 | 0.4106 | 0.0596 | 0.5675 | 0.0676 |
| CNNQA | 0.7126 | 0.0832 | 0.3981 | 0.0594 | 0.5553 | 0.0713 |
| LocalQA-net | 0.7040 | 0.0839 | 0.3620 | 0.0639 | 0.5330 | 0.0739 |
| MULTICOM-CLUSTER | 0.6543 | 0.0965 | 0.4006 | 0.0671 | 0.5275 | 0.0818 |
| JointQA-net | 0.6766 | 0.0954 | 0.3695 | 0.0648 | 0.5230 | 0.0801 |
| myprotein-me | 0.6498 | 0.0823 | 0.3875 | 0.0668 | 0.5187 | 0.0745 |
| MULTICOM-NOVEL | 0.6517 | 0.0949 | 0.3855 | 0.0623 | 0.5186 | 0.0786 |
| Wang_SVM | 0.6654 | 0.1011 | 0.3676 | 0.0828 | 0.5165 | 0.0920 |
| ProQ2-refine | 0.6640 | 0.0912 | 0.3530 | 0.0658 | 0.5085 | 0.0785 |
| ProQ2 | 0.6543 | 0.0862 | 0.3535 | 0.0570 | 0.5039 | 0.0716 |
| VoroMQA | 0.5787 | 0.1068 | 0.4010 | 0.0727 | 0.4899 | 0.0897 |
| RFMQA | 0.6275 | 0.0956 | 0.3485 | 0.0677 | 0.4880 | 0.0816 |
| Wang_deep_2 | 0.6395 | 0.1125 | 0.3115 | 0.0832 | 0.4755 | 0.0979 |
| Wang_deep_3 | 0.6338 | 0.1139 | 0.3050 | 0.0887 | 0.4694 | 0.1013 |
| Wang_deep_1 | 0.6208 | 0.1211 | 0.3071 | 0.0922 | 0.4640 | 0.1067 |

**Table 4:** The evaluation results (Corr. – Pearson’s correlation and loss) of global quality predictions of single-model QA predictors on stage 1 and stage 2 models of CASP 12 datasets.

|  | Stage1 |  | Stage2 |  | Average |  |
| --- | --- | --- | --- | --- | --- | --- |
| Feature | Corr | Loss | Corr | Loss | Corr | Loss |
| CNNQA | 0.7283 | 0.0627 | 0.6270 | 0.0854 | 0.6777 | 0.0740 |
| InteractQA-net | 0.7208 | 0.0613 | 0.6250 | 0.0844 | 0.6729 | 0.0728 |
| JointQA-net | 0.7124 | 0.0597 | 0.6230 | 0.0877 | 0.6677 | 0.0737 |
| Wang4 | 0.7105 | 0.0689 | 0.5965 | 0.1130 | 0.6535 | 0.0909 |
| LocalQA-net | 0.6928 | 0.0627 | 0.6112 | 0.0934 | 0.6520 | 0.0780 |
| MULTICOM-CLUSTER | 0.6958 | 0.0817 | 0.5895 | 0.0975 | 0.6427 | 0.0896 |
| SVMQA | 0.6540 | 0.0353 | 0.6227 | 0.0661 | 0.6383 | 0.0507 |
| ProQ3 | 0.6470 | 0.0524 | 0.6253 | 0.0710 | 0.6361 | 0.0617 |
| ProQ2 | 0.6677 | 0.0802 | 0.5981 | 0.0737 | 0.6329 | 0.0769 |
| Wang2 | 0.6713 | 0.0766 | 0.5329 | 0.1440 | 0.6021 | 0.1103 |
| VoroMQA | 0.6252 | 0.0798 | 0.5703 | 0.1027 | 0.5978 | 0.0912 |
| Wang1 | 0.4517 | 0.2002 | 0.2626 | 0.1486 | 0.3572 | 0.1744 |

### Influence of Rosetta energy terms and single-domain targets on the quality predictions

The performance of the three methods with and without Rosetta energies as input and trained on either the single-domain dataset or the full-length dataset was evaluated on Stage 1, Stage 2, and all the models of CASP11 and CASP12 test datasets and were visualized in the Figure 1. It is worth noting that each network with specific data input was fully tuned to the best performance on the validation dataset before being evaluated on the independent test dataset. As results shown in Figure 1, adding Rosetta energies improved the local quality prediction in most cases, with an average 1.26 improvement in ASE score. Training the network on the single-domain datasets also generally improved the performance over on the models of all targets (both single-domain and multi-domain targets, with an average 0.49 improvement on the CASP11 and CASP12 datasets in terms of ASE score.

**Figure 1:**
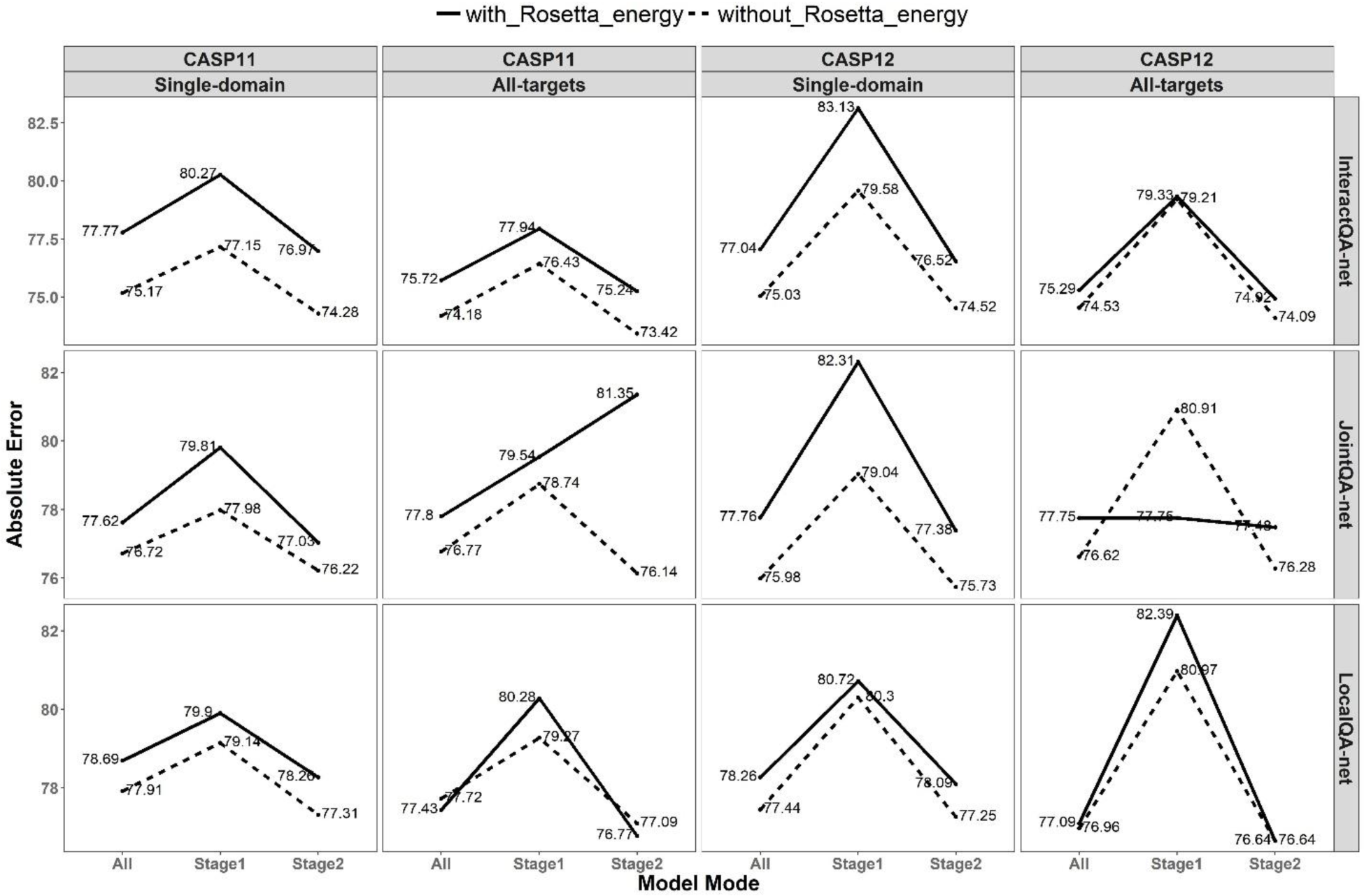
The comparison of local quality predictions of the method trained in different situations: (1) with Rosetta energy, (2) without Rosetta energh, (3) using only single-domain models in training, and (4) using both single-domain and multi-domain models (all models of all targets) in training.

### Case study of local quality predictions

**Figure 2** shows local quality predictions made by our method CNNQA for one model of target T0843 and one model of T0861 from the CASP12 experiment. The Figure 2(A) plots the real distance (gray) and predicted distance (green) at each residue position of the structural model of T0843, where the two curves overlap well at most positions. Figure 2(B) shows the superimposition of the native structure (gray) and the structural model(green). The average deviation between actual distances and predicted distances at all residue position in the model is 0.56 angstrom. The red highlighted regions have relatively large deviation (large errors) after the two structures being superimpose. Interestingly, these highlighted regions with a large distance deviation can be captured by the local quality prediction shown in Figure 2(A). Figure 2(C) and **2(D)** show the similar results for the model of T0861, where the average difference between real and predicted distance deviation is 0.67 angstrom.

**Figure 2:**
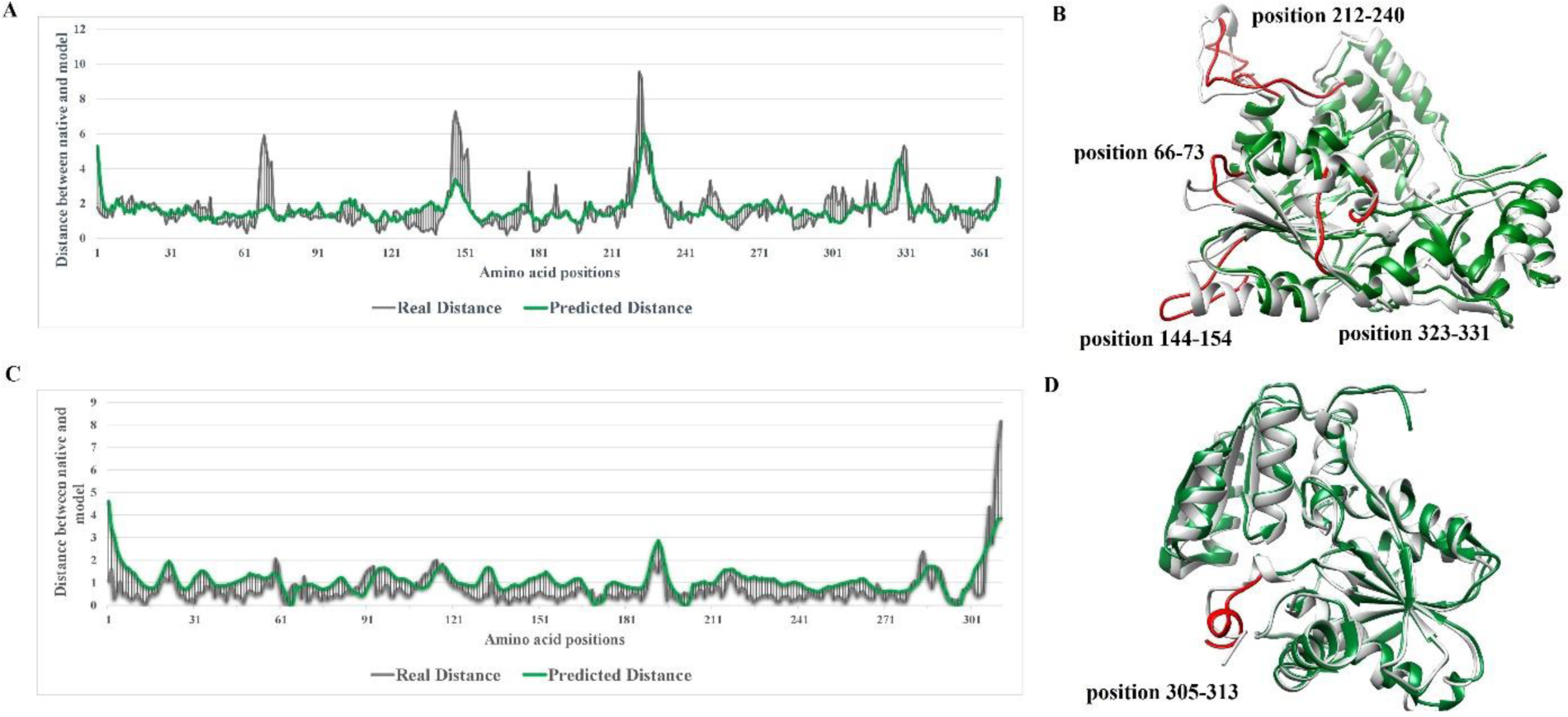
Residue-specific distance error predicted by our method (Green) and the real distance error between predicted model and native structure (Gray). (A) The distance error at each amino acid position in the predicted local quality and in the predicted model of T0843. (B) The superimposition of the predicted model (Green) of the model for target T0843 and its native structure (Gray). The red highlighted regions are the major deviation between predicted model and native structure, matching the large predicted deviation in the local quality prediction. (C) The distance error at each amino acid position in the predicted local quality and in the predicted model of T0861. (D) The superimposition of the predicted model (Green) of the mdoel for target T0861 and its native structure (Gray).

## Conclusion

In this work, we presented the novel 1D convolutional neural networks for predicting the quality of a single protein model. Instead of using fixed-size sliding windows to generate features for each residue, our network accepts the input of an entire protein model of arbitrary sequence length and therefore it can access the global structural information that informs the quality of a position of residue. We also designed a new training pipeline to integrate local and global quality prediction together, which improved the accuracy of global quality prediction. Overall, our methods performed comparably to the state-of-the-art methods in the past CASP11 and CASP12 experiments. The results demonstrate that 1D deep convolutional neural networks are promising techniques for protein model quality assessment. In the near future, we will design more advanced deep learning architectures to further advance protein model quality prediction.

## Methods

### Datasets

The dataset for training and validation was downloaded from the 8th, 9th and 10th Critical Assessments of Structure Prediction (CASP) experiments (http://predictioncenter.org/), consisting of the models for 322 protein targets whose native structures were officially released. The targets with multiple-domains were removed from dataset because using only single-domain models to train the methods worked better (see the Results and Discussions Section for details), and the remaining protein models for single-domain targets were used for training and validation, leading to 48,574 structural models for 236 single-domain targets. Specifically, the final dataset contains 15,022 models of 82 CASP8 targets, 19,926 models of 87 CASP9 targets, and 13,626 models of 67 CASP10 targets.

The 236 targets were randomly split into the two sets according to the 80% - 20% ratio. The models of 80% targets were used for training and the rest for validation and parameter tuning. Specifically, the final training dataset contains 38,832 models and the validation dataset contains 9,742 models. The independent test datasets include the models of all the single-domain and multi-domain targets of CASP11 and CASP12 experiments. Specifically, 14,076 models of 84 CASP11 targets and 6,008 models from 40 CASP12 targets whose native structure were released to date were included into the test dataset.

The true local and global quality scores of the models in the datasets above were obtained by comparing them with the corresponding native structures. The local quality and global scores predicted by other CASP QA methods for the models were downloaded directly from CASP data repository (http://predictioncenter.org/) for comparison with our methods.

### Feature Extraction

Our one-dimensional deep convolutional networks (1DCNN) take the following residue-wise raw features and several global features as input, which include (1) amino acid encoding of each residue, (2) position specific scoring matrix (PSSM) profile of each residue derived from the multiple sequence alignment of the protein, (3) predicted secondary structure of each residue, (4) predicted solvent accessibility of each residue, (5) predicted disorder state of each residue, (6) the agreement between the secondary structure of each residue in the model and the predicted one and, (7) the agreement of solvent accessibility of each residue in the model and the predicted one, (8) Rosetta energies of each residue as in the ProQ3^15^, which is calculated from Van der Waals, side-chains, Hydrogen bonds, and Backbone information, and (9) six global knowledge-based potentials or features of the entire model produced by ModelEvaluator^20^, Dope^18^, RWplus^30^, Qprob^14^, GOAP^38^, and Surface score. The amino acid encoding is a vector of 20 binary numbers where the value at the index of the residue index is labeled as 1, otherwise as 0. The PSSM profile is generated by PSI-BLAST^39^ searching the sequence against ‘nr90’ sequence database. SSPro^40^ was run to generate the predicted secondary structure and solvent accessibility for each residue in the model. The disorder state of each residue was predicted by PreDisorder^41^. The features of a model of L residues are stored in a vector of length L. Each element of the vector contains all the local features of each residue as well as several global features, which is the input for the deep convolutional neural network.

We used LGA structural alignment tool^42^ to measure the local residue-wise distance error and global structural similarity score between models and their native structures. The local distance error is defined by the distance deviation of each residue in a model and in the native structure after superimposing them together, while the global similarity score is defined by the GDT-TS score^43^ – the average percent of residues in the model that are close to their positions in the native structure according to several thresholds. We used a function S-function 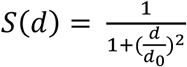 applied in the previous studies ^8,10^ to scale the local distance deviation of residues into the range of [0,1], where *d* is the distance deviation of a residue between model and native structure, and *d*_0_ is set to 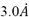. Lower a distance, higher is the *S* score. *d* and S can be converted back and forth.

### Deep convolutional neural network for protein model quality prediction

We designed three architectures of deep convolutional neural networks (CNN) for predicting the residue-wise local quality of a protein model and investigating the effect of global quality prediction on the local quality prediction. Our first network (LocalQA-net) is designed for local quality prediction using 1D convolutional neural networks, as shown in Figure 3(A). The network has one input layer for each protein structure of any length, multiple hidden convolutional layers and one output layer to predict final residual qualities of the same size. In the hidden convolutional layers, the “rectified-linear unit (ReLU)” activation function^44^ and batch normalization^45^ were applied during training. Our second network (InteractQA-net) consists of two sub-networks for local quality and global quality prediction separately and a common convolutional sub-network of extracting features from the input layer that are shared by the former two, as shown in Figure 3(B). On top of the common convolutional sub-network, the sub-network for predicting local quality, referred to as LocalQA-net, has one convolutional output layer with a sigmoid activation function to predict the local quality score for each residue in a model, resulting in L scores for a model of length L. The sub-network for the global quality prediction, referred to as GlobalQA-net, shares the same common network as LocalQA-net, followed by one K-max pooling layer^46^ (default K=30), one standard fully connected layer (default 50 hidden nodes), and one single output node to predict the global quality score of an input model. Given a protein model, the InteractQA-net first optimized the weights of LocalQA-net and the common sub-network based on local quality scores. Then the shared weights in the convolutional layers were transferred to GlobalQA-net and both the shared weights and the weights of GlobalQA-net were optimized by global quality scores. After the weights were updated by training on GlobalQA-net, the shared weights were transferred back to LocalQA-net for further optimization. These steps iterated until training converged or the maximum number of iterations was reached. The network was optimized by the Nesterov Adam (nadam)^47^ method with Mean Square Error (MSE) as loss function. In order to optimize the performance, we adjusted three main hyper-parameters of convolutional layers during training, including (1) the depth of the network (from 5 to 10), (2) the filter size of the filters in the convolutional layer (from 5 to 10), and (3) the number of filters in each convolutional layer (from 5 to 20). The c (referred as ASE)^48^, a standard measure used in CASP to assess the accuracy of local quality prediction, for each parameter setting on the validation was calculated. ASE is the averaged absolute difference of predicted quality score and real quality score of each residue in a model. ASE is defined as 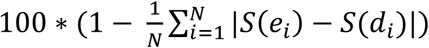, which is (1 - the average difference of predicted residue quality (*S*(*d*_*i*_)) and real residue quality (*S(e*_*i*_))) times 100. The higher ASE score, more accurate is the local quality prediction. The parameter setting yielding higher ASE was preferred. Each convolutional layer applies the batch-normalization and uses the rectified-linear unit (ReLU) activation function to convert its activation into its output.

**Figure 3:**
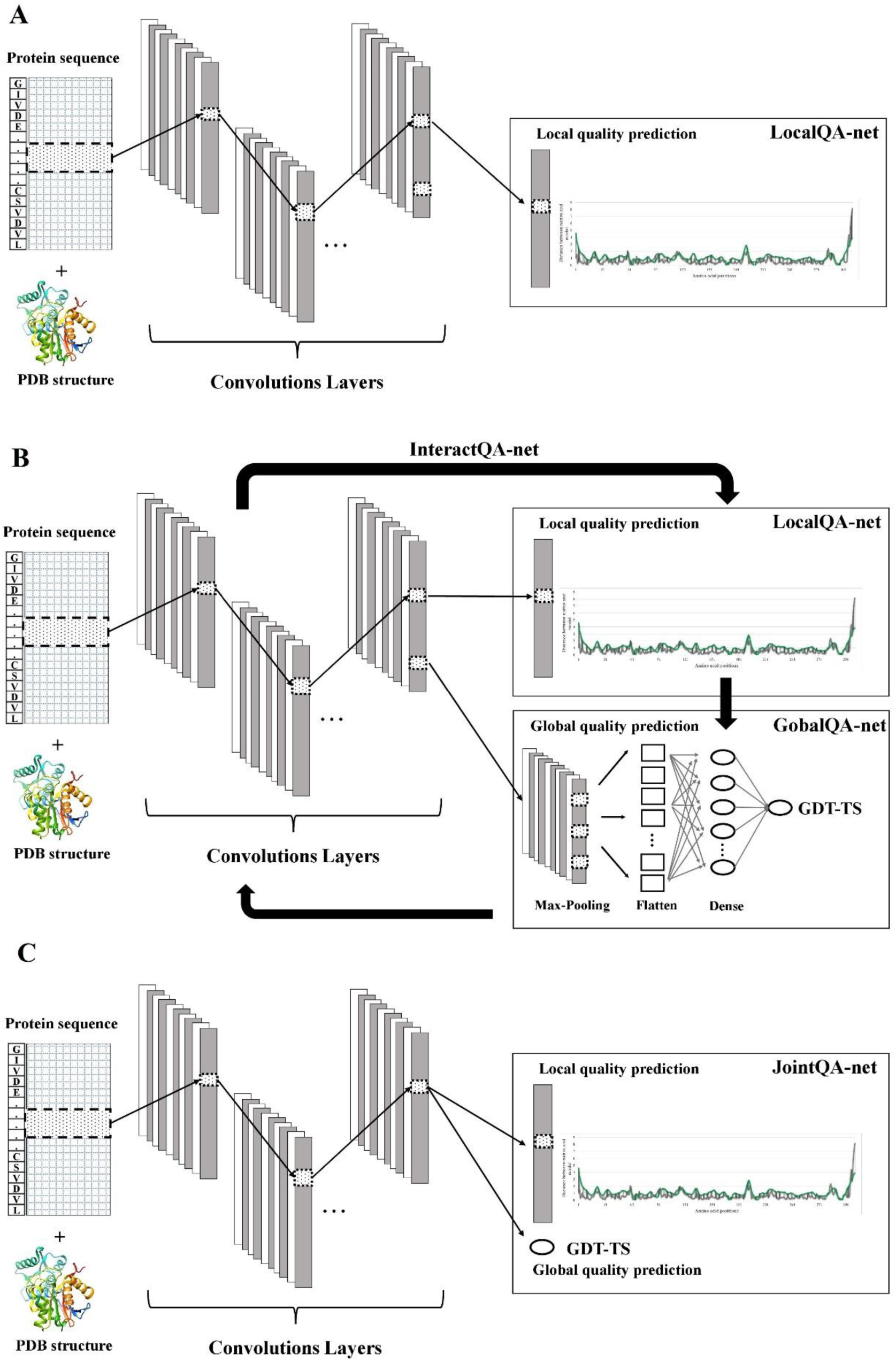
The architecture of 1D deep convolutional neural network for protein model quality prediction. **(A).** The network (LocalQA-net) accepts the raw features of models of proteins of variable sequence length (L) as input, and transforms the features into higher-level hidden features by 5 hidden layers of convolutions. Each convolutional layer applies 5 filters to windows of previous layers to generate L hidden features. The window size for each filter is set to 6. The last output layer adds one convolutional layer with one filter to generate the output of length L representing the local quality for each of L residues. **(B)** The network (InteractQA-net) contains a common sub-network for extracting features from the input layer by convolution and two sub-networks for local quality (LocalQA-net) and global quality (GlobalQA-net) predictions separately. The network accepts the raw features of models of proteins of variable sequence length (L) as input, and transforms the features into higher-level hidden features by 10 hidden layers of convolutions. Each convolutional layer applies 10 filters to windows of previous layers to generate L hidden features. The window size for each filter is set to 15. LocalQA-net adds one convolutional layer with one filter at the top of the common sub-network to generate the output of length L representing the local quality for each of L residues. GlobalQA-net uses one 30-max pooling layer to select 30 maximum values from the output of each filer in the last layer of the common sub-network as features, which are joined together into one vector by a flatten layer. The flatten layer is fully connected to a hidden layer whose output is used by a single output node to predict the global quality score. LocalQA-net and GlobalQA-net are trained by local quality scores and global quality scores alternately. **(C)** JointQA-net accepts the features of protein models of variable sequence length (L) as input and predicts the L local quality scores and one global quality score simultaneously. The weights in the network are optimized by both local and global quality scores at the same time.

In addition to the architecture above, we also designed another architecture called ‘JointQA-net’ to integrate global quality prediction with local quality prediction, as shown in Figure 3(C). The common sub-network and the sub-network for local quality prediction in JointQA-net are the same as InteractQA-net. But JointQA-net has a much simpler sub-network for global quality prediction, which has only one single output node to predict global quality scores. Moreover, instead of alternately training networks using local quality scores and global similarity scores as ‘InteractQA-net’, JointQA-net predicts both quality scores simultaneously in its output layer in order to optimize all the weights in the network at the same time. Finally, in order to evaluate the effectiveness of incorporating global quality predictions into local quality learning, both InteractQA-net and JointQA-net were compared to the basic network of local quality prediction - LocalQA-net – whose weights were not adjusted according to global quality scores but according to local quality scores only.

### Evaluation and Benchmarking

We evaluated both local and global quality predictions of our deep learning methods on two sub-sets (1^*st*^ stage and 2^*nd*^ stage) of CASP11 and CASP12 datasets, respectively. The local quality predictions were evaluated based on the ASE score^48^. The global quality predictions were evaluated in terms of (1) Pearson’s correlation between predicted global scores of the models of a target and the real global quality scores of the models of the target, and (2) the average loss. The loss is the difference between the real quality score of the no. 1 model selected according to predicted quality scores for a target and the quality score of the real best model of the target. The average loss evaluates the capability of a method to select good models. A loss 0 means the predicted global quality scores can always rank the real best model as no. 1.

We evaluated the performance of our three local quality predictors (InteractQA-net, JointQA and LocalQA-net) on the test datasets and compared them with other QA predictors that participated in CASP 11 and CASP 12. The predictions of CASP QA predictors were directly downloaded from CASP repository (http://predictioncenter.org/). In order to evaluate the performance of our global quality predictions, we converted the local quality prediction made by InteractQA-net, JointQA and LocalQA-net into global quality scores by averaging the local quality predictions of residues directly using function 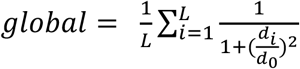. Besides, an ensemble of our three predictors called CNNQA, which uses the average output of the three predictions as its prediction, was evaluated.

## Acknowledgements

The work was partially supported by an NIH grant (R01GM093123) and two NSF grants (DBI1759934 & IIS1763246) to JC.

## Affiliations

Department of Electrical Engineering and Computer Science, University of Missouri, Columbia, Missouri, 65211, USA. Jie Hou & Jianlin Cheng

Informatics Institute, University of Missouri, Columbia, Missouri, 65211, USA. Jianlin Cheng Department of Computer Science, Pacific Lutheran University, WA 98447, USA. Renzhi Cao

## Author Contributions

JC and JH conceived and designed the experiment. JH implemented the method. JH and RC performed the analysis. All the authors wrote the manuscript.

## Corresponding author

Correspondence to Jianlin Cheng.

## Competing interests

The authors declare no competing interests.

